# The trimeric Shu complex in *C. elegans* is an ATPase that remodels RAD51 filaments in the homologous recombination-associated DNA damage response

**DOI:** 10.1101/2025.11.17.685403

**Authors:** Sam S. H. Chu, Guangxin Xing, Hong Ling

**Author notes:** Correspondence should be addressed to H. L.

## Abstract

Homologous recombination (HR) is critical to error-free lesion bypass in the DNA damage response. This error-free lesion tolerance pathway is initiated by the RAD51 recombinase, which forms nucleoprotein filaments on single-stranded DNA (ssDNA) to facilitate template-directed repair using homologous sister chromatids. RAD51 filaments are tightly regulated by RAD51 mediator proteins. Among these, the Shu complexes facilitate HR-directed DNA damage tolerance and are evolutionarily conserved from yeast to humans. The *Caenorhabditis elegans* Shu complex is a heterotrimer consisting of three protein subunits: RFS1, RIP1, and SWS1. However, the biochemical properties of this trimeric complex remain unclear. Here, we report the biochemical characterization of the *C*. *elegans* Shu complex and interactions with DNA, ATP, and Rad51 filaments. We first revealed that the Shu trimer preferentially binds DNA with an exposed 5′ end, particularly favoring a fork-shaped double-stranded DNA (dsDNA). Then, we found that the trimer binds to ATP and exhibits DNA-dependent ATPase activity. Through site-specific mutagenesis, we identified the catalytic residues in the RFS1 domain and validated the ATPase activity. Using fluorescence-based assays, we further demonstrated that the Shu trimer remodels RAD51 filaments in an ATP-hydrolysis-dependent manner and stabilizes the filaments in an ATP-binding-dependent manner. These findings provide key mechanistic insights into how the *C*. *elegans* Shu complex regulates RAD51 filaments, priming them for downstream HR-mediated DNA repair processes.

## Introduction

Genomic integrity and cell survival are constantly challenged by endogenous and environmental DNA-damaging agents, causing numerous DNA lesions daily^1–4^. In the S and G_2_ phases of the cell cycle, persistent replication-blocking DNA lesions require the homologous recombination (HR) pathway as part of the error-free DNA damage tolerance mechanism^5^. HR-associated DNA lesion bypass is considered error-free because sister chromatids serve as repair templates, ensuring accurate DNA replication and minimizing mutations that could lead to genome instability and cancers^5,6^. A key player in HR is the highly conserved eukaryotic RAD51 recombinase, which forms filaments on resected single-stranded DNA (ssDNA)^6,7^. RAD51 filaments, in conjunction with RAD51 mediator proteins, facilitate homology search, strand invasion, and strand exchange with the homologous sister chromatids during DNA lesion bypass^5,6,8,9^. The Shu complexes, conserved from yeast to humans, are RAD51 mediators involved in the DNA damage response^5,10–14^. Previous studies in yeast and human systems have established the Shu complex as a key regulator of HR, promoting RAD51 filament formation and suppressing the dismantling activity of anti-recombinases, such as Srs2 in yeast and FIGNL1 in humans^10,14–17^. Complementing these systems, *Caenorhabditis elegans* (*C*. *elegans*) has emerged as a powerful and genetically tractable model for dissecting Shu complex function^13^. The highly ordered germline allows for direct visualization of meiotic progression, while programmed DNA double-strand breaks (DSBs), induced by the conserved enzyme SPO11, are repaired primarily by HR-mediated damage responses^13,18,19^. Moreover, the self-fertilizing reproductive system enables efficient genetic screening, with meiotic defects inferred from progeny phenotypes, such as increased male frequency or embryonic lethality^13,20^. Thus, biochemical analysis of the *C*. *elegans* Shu complex is essential to uncover the molecular mechanisms underlying these phenotypes and to directly define the role of the complex in RAD51 filament dynamics.

The *C*. *elegans* Shu complex (cShu) consists of three protein subunits: RFS1, RIP1, and SWS1. RFS1 and RIP1 have been identified as RAD51 paralogs and contain Walker A and B motifs, which are conserved for ATP binding and hydrolysis, respectively^12^. SWS1 was later identified as a binding partner of the RFS1-RIP1 dimer through yeast two-hybrid experiments^13^ and contains a conserved SWIM domain, a characteristic feature of Shu homologs^14^. Biochemically, the function of the trimeric cShu is largely uncharacterized. Previous studies demonstrated that the RFS1-RIP1 dimer regulates RAD51 filament assembly, conformation, and stability in an ATP-binding-dependent manner^12,21–23^. Comparatively, the complete yeast Shu complex (Csm2-Psy3-Shu1-Shu2) exhibits DNA-dependent ATPase activity that is essential for modulating RAD51 filament properties^24^. Similarly, in humans, the hSWS1-SWSAP1 complex binds RAD51 to form interspersed filaments^25^ and exhibits DNA-stimulated ATPase activity^10,26^. More recently, ATP binding and hydrolysis by the human Shu complex were shown to play distinct roles in RAD51 filament modulation^26^. The SWIM domain-containing subunit has been shown to be essential for ATPase function in both yeast and human Shu complexes, and this activity is critical for RAD51 filament remodeling^10,24,26^. Furthermore, disruptions to the conserved SWIM domain impair DNA damage tolerance in both organisms^14^, underscoring the importance of Shu complex ATPase activity in HR-mediated DNA lesion bypass. Since it remains unclear whether the RAD51 filament-modulating activity of the complete *C*. *elegans* Shu complex is strictly ATP-binding-dependent, detailed biochemical characterization of the trimer is essential for elucidating the mechanistic role of the complex as a RAD51 mediator.

In this study, we biochemically characterized the *C*. *elegans* Shu complex, RFS1-RIP1-SWS1, to elucidate the function of the complex in HR. We demonstrated that cShu preferentially binds DNA with an exposed 5′ end. We characterized the ATP-binding and hydrolysis activities of cShu and found that the ATPase activity is stimulated by DNA. Finally, we showed that cShu modulates RAD51-ssDNA filament properties, specifically altering filament conformation in an ATP-hydrolysis-dependent manner, while stabilizing filaments in an ATP-binding-dependent manner.

## Materials and Methods

### Generation of recombinant bacmids for insect cell protein expression

The plasmids, pUC57-SWS1, pUC57-RFS1, and pUC57-RIP1, encoding each of the respective *C*. *elegans* Shu complex genes, were purchased from Biobasic. We built all the plasmid constructs using the polymerase incomplete primer extension (PIPE) method^27^. The protocol for generating recombinant bacmids was adapted from the MultiBac™ Kit (Geneva Biotech) manual, with modifications. Briefly, *RFS1*, *RIP1*, and *SWS1* were cloned into a modified pACEBac1 vector for co-expression of a TEV-cleavable 6×His-tagged RFS1 together with native RIP1 and SWS1. The K56A (Walker A) and E138A (Walker B) mutants of RFS1 were generated for functional validation. Transfer vectors of pACEBac1-6×His-TEV-RFS1-RIP1-SWS1 and the Walker-motif mutants were transformed into *E*. *coli* competent cells harboring the DH10EMBacY bacmid. Tn7 transposition integrated the genes of interest into the recipient baculovirus. Recombinant bacmids carrying the integrated multigene cassettes were selected by blue/white screening and gentamicin resistance. Isolated recombinant bacmids with genes of interest were verified using PCR analysis, as outlined in the Bac-to-Bac^®^ Baculovirus Expression System (Invitrogen) manual.

### Expression and purification of C. elegans Shu complex and RAD51

The RFS1-RIP1-SWS1 trimer and the Walker-motif mutants were expressed in insect cells as described^28^, with modifications. Prior to transfection, Sf9 cells were maintained in Gibco Sf900-II SFM (Thermo Fisher Scientific) at a density of 1.0 to 4.0 × 10^6^ cells/ml with a viability of 95% or higher. EMBacY-6×His-RFS1-RIP1-SWS1 recombinant bacmids were heat-sterilized at 55 °C for 1 hour. The bacmid-polyethyleneimine mixtures, incubated at room temperature for 30 min, were added dropwise to the diluted Sf9 cell culture. The transfected Sf9 cell cultures were incubated at 27 °C with shaking. Five days post-infection, the media containing recombinant baculovirus were collected and mixed with fetal bovine serum.

High Five cells were maintained in I-MAX insect cell culture media (Wisent Bio Products) at a density of 1.0 to 4.0 × 10^6^ cells/ml with a viability of 95% or higher, and infected with each recombinant baculovirus. Cell cultures were harvested by centrifugation three days post-infection or at 85% cell viability. Cell paste was resuspended and washed with 1X phosphate-buffered saline (PBS) buffer.

Cell paste was lysed in a lysis buffer (50 mM Tris-HCl, pH 7.5, 0.5 M NaCl, 5% glycerol, 1 mM PMSF, 1 mM benzamidine, 1 mM E-64, 1 mM bestatin, 1 mM pepstatin A, and 5 mM βMe). Clarified cell lysate was purified using a Ni-HiTrap FF column (GE Healthcare), further purified by gel filtration chromatography using a Superdex 200 10/300 GL column (GE Healthcare), and stored in a storage buffer (30 mM Tris-HCl, pH 8.0, 0.25 M NaCl, and 2 mM DTT). The purified proteins were of high purity (>95%) and homogeneity as determined by Bis-Tris-SDS-PAGE.

*RFS1* and *RIP1* were cloned into pMCSG9 and a pMCSG7 derivative, respectively, to co-express RFS1 with a His-MBP tag and RIP1 with a His-MOCR tag (both TEV-cleavable). The plasmid encoding *C*. *elegans* RAD51 was kindly provided by Dr. Simon J. Boulton of Clare Hall Laboratory at The Francis Crick Institute, which expresses His-SUMO-RAD51 (Ulp1-cleavable). The plasmid was transformed into *E*. *coli* BL21 (DE3) codon plus strain for overexpression. Cell cultures were grown to an OD_600_ of 0.6–0.7, induced at 16 °C for 16 hours, and harvested by centrifugation and stored at -80 °C until protein purification at 4 °C.

Cell paste was lysed in a lysis buffer (50 mM Tris-HCl, pH 7.5, 0.5 M NaCl, 5% glycerol, 1 mM PMSF, 1 mM benzamidine, 1 mM E-64, 1 mM bestatin, 1 mM pepstatin A, and 5 mM βMe). Clarified cell lysates were purified using a Ni-HiTrap FF column (GE Healthcare). Tag-cleaved proteins were then purified with ion exchange chromatography (GE Healthcare Q FF column). The protein of interest was eluted by a linear gradient of NaCl. RFS1-RIP1 dimer was further purified using gel filtration chromatography on a Superdex 200 10/300 GL column (GE Healthcare). RFS1-RIP1 dimer and RAD51 were stored in storage buffer (30 mM Tris-HCl, pH 8.0, 0.25 M NaCl, and 2 mM DTT). The purified proteins were of high purity (>95%) and homogeneity as determined by Bis-Tris-SDS-PAGE.

Protein concentrations were quantified using the Bio-Rad Protein Assay Dye Reagent in a colorimetric assay. Final protein concentrations were determined from a standard curve generated using bovine serum albumin (BSA) (New England Biolabs).

### Electrophoretic mobility shift assays (EMSAs) for DNA binding

DNA-binding reactions were carried out with increasing concentrations of RFS1-RIP1-SWS1 trimer proteins or RFS1-RIP1 dimer and 0.05 μM DNA substrate labeled with 6-carboxyfluorescein (6-FAM). The protein-DNA mixtures were resolved on native polyacrylamide gels and imaged using the ChemiDoc MP Imaging System (Bio-Rad). Assays were performed in triplicate.

### Fluorescence polarization assays (FPAs) for DNA binding

FPAs for DNA-binding analysis of RFS1-RIP1-SWS1 trimer proteins and RFS1-RIP1 dimer were conducted as described previously^24,26,29,30^. Fluorescence polarization measurements were recorded using a VICTOR^3^ V 1420 Multilabel Counter (PerkinElmer) with an integrated polarizer.

Reactions were carried out at 20 °C. Each reaction mixture contained 5 nM 6-FAM-labeled DNA substrates with increasing concentrations of proteins. Changes in fluorescence polarization were obtained by subtracting the measurement of DNA alone from each data point. Triplicate data were fitted using a one-site binding model in PRISM 5 (GraphPad) to a quadratic equation **(1)** to determine the *K*_d_ values. DNA concentrations in molecules were converted to concentrations in nucleotides for curve fitting.

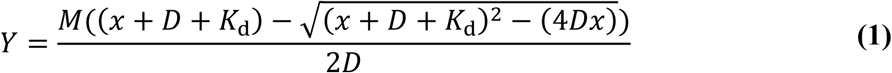

### Analysis of protein-ATP interactions with a fluorescent ATP analog

TNP-ATP spectra, for ATP-binding analysis of RFS1-RIP1-SWS1 trimer proteins and RFS1-RIP1 dimer, were collected under conditions described previously^24,26,31^. Experiments were performed using a Synergy H1 Hybrid Multi-Mode Microplate Reader (BioTek Instruments) at 20 °C. For the TNP-ATP saturation assays, emission readings at 540 nm were collected for 2 μM protein with increasing TNP-ATP concentrations. The correction factors for the inner filter effect were established and applied to data points above 10 μM TNP-ATP as previously described^32–34^. Triplicate data were fitted in PRISM 5 (GraphPad) by non-linear regression with a one-site specific binding model to determine *K_d_^TNP-ATP^*, using equation **(2)**.

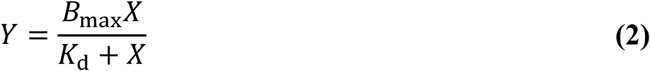

For the competitive binding curve, 2 μM cShu with fixed 2.5 μM TNP-ATP was titrated with increasing concentrations of ATP. Triplicate data were fitted in PRISM 5 (GraphPad) by non-linear regression with a one-site competitive binding model to determine the *K*_i_^ATP^, using equations **(3)** and **(4)**.

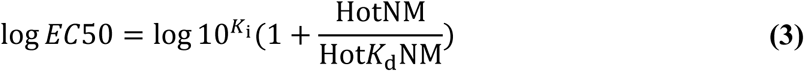

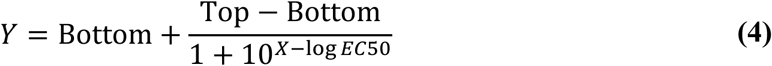

The competitive binding data set was then used to derive the corresponding *K_d_^ATP^* parameter, using equations **(5)** and **(6)**^35^.

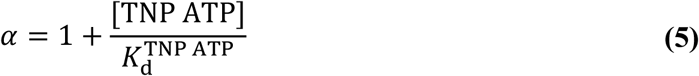

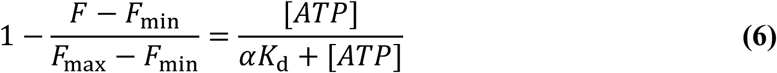

### ATPase activity assays

ATPase activity assays of RFS1-RIP1-SWS1 trimer proteins and RFS1-RIP1 dimer were performed as described^24,26,36,37^, with slight modifications. The assays were performed with 1 μM proteins in the presence and absence of 10 μM DNA at 20 °C.

Steady-state ATPase kinetic assays with cShu were performed with/without DNA (1 μM), 1 μM protein, and increasing concentrations of ATP. Triplicate data sets were used to calculate the kinetic parameters *K*_m_, *V*_max_, and *k*_cat_ with a non-linear regression fit of the nanomoles Pi released/min to the Michaelis–Menten equation **(7)**.

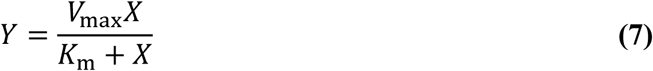

### Nuclease protection assays

DNase I digestion assays were performed as described^24,26^. The 5′- or 3′-6-FAM-labeled ssDNA substrates (0.25 μM) were mixed with cShu proteins. After 30 min incubation at 24 °C, 1 mM CaCl_2_ and 2 Units bovine pancreatic DNase I (New England Biolabs) were added to protein-DNA mixtures at 37 °C. For reactions with *C*. *elegans* RAD51 filaments, RAD51-ssDNA was incubated for 10 min at 24 °C prior to the addition of cShu proteins, and further incubated for 30 min. Reaction mixtures were stopped by the addition of a mixture containing EDTA and formamide, and subsequently resolved on denaturing polyacrylamide urea gels. Gel images were imaged using the ChemiDoc MP Imaging System (Bio-Rad). Assays were performed in triplicate.

### Fluorescence assays of C. elegans Shu complex-Rad51 filaments

Fluorescence assays of RAD51 filaments with RFS1-RIP1-SWS1 trimer proteins and RFS1-RIP1 dimer were performed as described^24,26^. Samples were excited at 490 nm, and fluorescence measurements were collected at the emission wavelength of 522 nm at 20 °C, using a Synergy H1 Hybrid Multi-Mode Microplate Reader (BioTek Instruments). Reactions consisted of *C*. *elegans* RAD51-ssDNA filaments (1 μM RAD51 and 15 nM of 5′- or 3′-6-FAM-labeled poly(dT)_39_ ssDNA), 2 mM ATP, and the *C*. *elegans* Shu proteins (50 nM). For reactions involving adenine nucleotide preincubation of cShu, the ATP, ADP, or AMP-PNP were incubated with the RFS1-RIP1-SWS1 trimer. Adenine nucleotide-preincubated cShu or each individual adenine nucleotide type (negative controls) was mixed with ATP-bound RAD51 filaments after the initial 10-minute incubation. In competition assays, a 100-fold excess of unlabeled poly(dT)_39_ ssDNA (1.5 μM) was mixed with the RAD51 filament/cShu protein mixture after the initial 10-minute incubation. Fluorescence readings of labeled ssDNA were used for background subtraction of all data sets. Fluorescence of the RAD51-ssDNA filament alone was measured for 60 seconds at 10-second intervals, with the first measurement defined as a common 0-second time point for all reactions. Subsequent RAD51-ssDNA fluorescence intensity measurements at each time point were used to normalize the corresponding time points for all other data sets.

## Results

### DNA-binding preferences of cShu

As a mediator of RAD51 filaments, characterizing the DNA-binding properties of the trimeric cShu provides valuable insights into the molecular role of the complex in HR. To systematically investigate the DNA-binding preferences of RFS1-RIP1-SWS1, we conducted DNA-binding assays using fluorescein-labeled DNA substrates of varying lengths and end types (**Table 1**). The trimeric cShu and the Walker-motif mutant variants were purified to near homogeneity with the correct oligomeric size (**Figure 1A & 1B**). We first tested DNA binding using EMSAs in two-layer native gels: a 5% PAGE layer at the top to allow large DNA-protein complexes to enter and a 15% PAGE layer at the bottom to prevent the free DNA from running out. The results showed that cShu physically binds both ssDNA and fork-shaped dsDNA substrates in a concentration-dependent manner but does not interact with the blunt-ended dsDNA substrate (**Figure 1C**). This suggests a binding preference for ssDNA over blunt-ended dsDNA.

**Figure 1.**
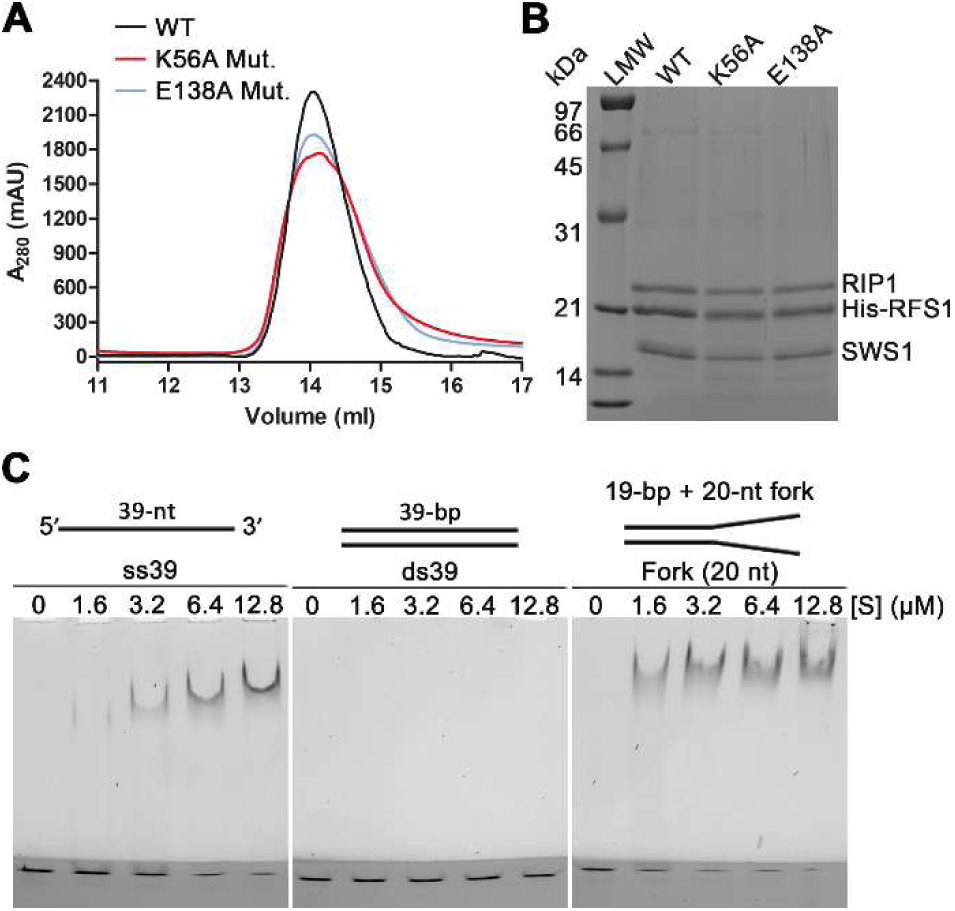
DNA-binding activity of purified trimeric *C*. *elegans* Shu complex (cShu). (**A**) Gel filtration and (**B**) SDS-PAGE of purified trimeric cShu (RFS1-RIP1-SWS1) wild type (WT) and mutants (K56A and E138A in RFS1). “LMW” stands for low-molecular-weight protein ladder. (**C**) Electrophoretic mobility shift assay (EMSA) for WT RFS1-RIP1-SWS1-DNA interactions. Increasing concentrations of RFS1-RIP1-SWS1 ([S]) were mixed with 0.05 μM DNA substrates (ss39, ds39, Fork (20 nt)). Protein-DNA mixtures were resolved by two-layer PAGE gels: 5% native polyacrylamide at the top and 15% at the bottom (dark layer). The number in parentheses indicates the number of nucleotides in the single-stranded parts of the “Fork (20 nt)” dsDNA in **C**. Assays were performed in triplicate with comparable results.

**Table 1.**
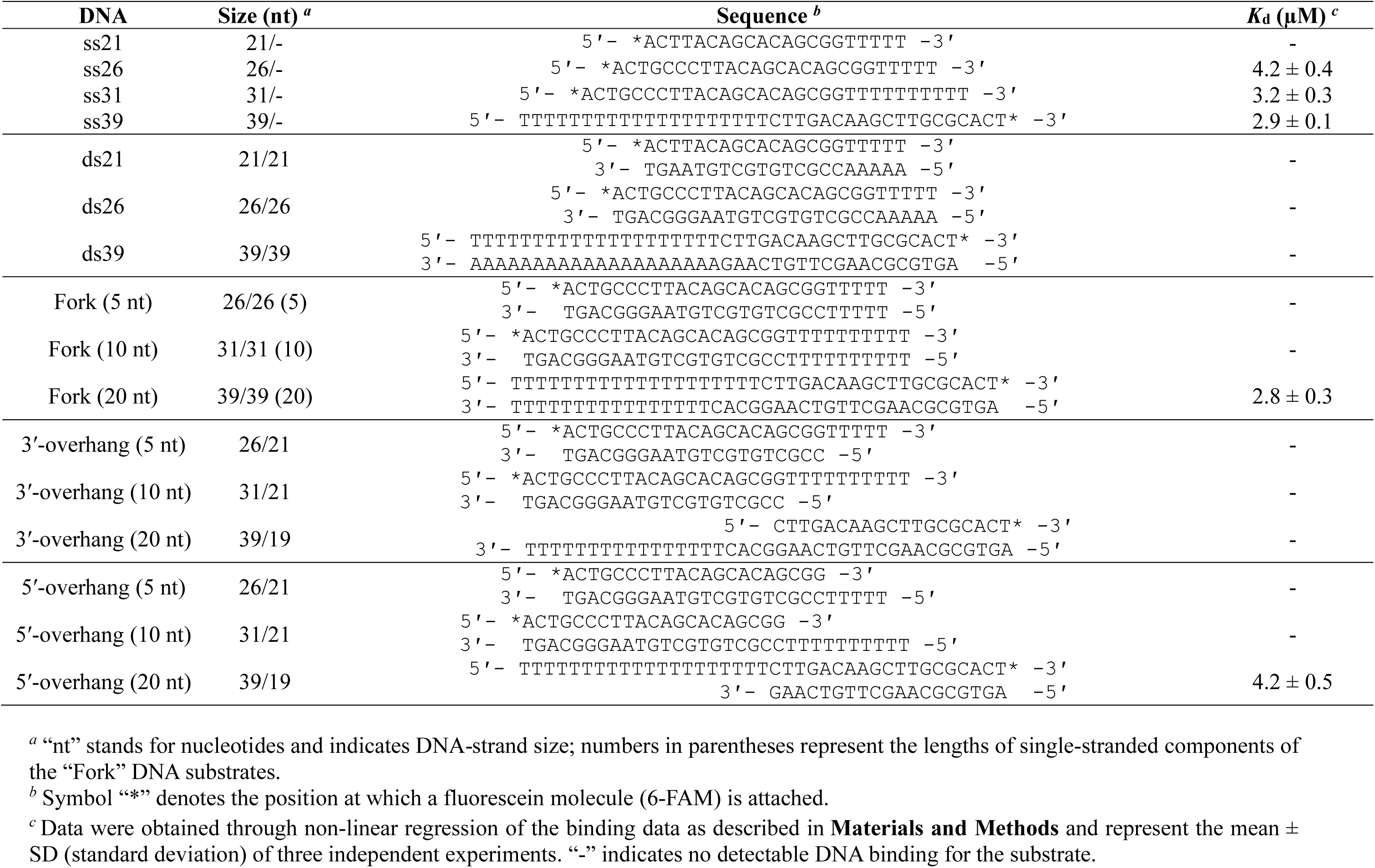
DNA-binding affinities of trimeric cShu.

We then conducted fluorescence polarization assays (FPAs) to determine the binding affinities of cShu with different DNA substrates (**Table 1, Figure S1A–S1C**). For ssDNA, cShu bound substrates ranging from 26 to 39 nucleotides (nt) in length, but not to the 21-mer substrate. (**Table 1, Figure S1A**). The *K*_d_ values for the ssDNA substrates ranged from 2.89 to 4.19 μM, averaging ∼3.44 μM (**Table 1**). Overall, the data suggest that cShu binds ssDNA with a minimum length longer than 21 nt. Consistent with the EMSA results (**Figure 1C**), cShu did not bind the blunt-ended dsDNA substrates (**Table 1, Figure S1A**).

Next, we examined the influence of end types on the DNA binding of cShu. In yeast, the Shu complex preferentially binds fork-shaped dsDNA substrates^24,38^ and exhibits strand-specificity^39^. It is likely that cShu exhibits a similar substrate preference and recognition mechanism. Hence, we first tested fork-shaped substrates with 5-, 10-, and 20-nt arms. cShu only bound the fork-shaped substrate with 20-nt arms (**Table 1, Figure S1B**), suggesting a minimum ssDNA arm length requirement. Based on the ssDNA-binding results, a length of 26 nt was necessary for the cShu-ssDNA interaction (**Table 1**). Since the fork-shaped substrate contains only 20-nt arms, the ssDNA/dsDNA junction likely contributes to binding.

We subsequently tested the strand specificity of cShu using 5′- and 3′-overhang dsDNA substrates with 5-, 10-, and 20-nt overhangs. Interestingly, cShu only bound the 5′-overhang (20 nt) substrate, but not the 3′-overhang counterpart (**Table 1, Figure S1C**), suggesting a preference for exposed 5′ ends. The *K*_d_ values for the fork-shaped and 5′-overhang dsDNA substrates were comparable to those of ssDNA (**Table 1**). In addition, the lack of binding for 5′-overhang substrates with overhangs shorter than 20 nt suggests that cShu requires a sufficiently long ssDNA segment with an exposed 5′ end for DNA binding, with the ssDNA/dsDNA junction contributing to the interaction. These findings align with the lagging-strand-specific binding preference of the yeast Shu complex^39^.

We further analyzed the DNA-binding preferences of cShu in the presence of ATP or ADP to test if adenine nucleotide binding influences DNA binding (**Table 2, Figure S1D – S1F**). The binding assays showed that ATP increased DNA-binding affinities for fork-shaped, 5′-overhang, and ssDNA substrates by ∼14-fold, ∼6-fold, and ∼3-fold, respectively (**Table 2**). In contrast, ADP increased fork-shaped dsDNA-binding affinity by ∼6-fold but did not affect the binding of 5′-overhang dsDNA and ssDNA (**Table 2**). The lack of binding to blunt-ended dsDNA and 3′-overhang substrates persisted regardless of adenine nucleotide presence (**Table 2, Figure S1E & S1F**). These results suggest that adenine nucleotide binding may regulate the DNA binding of cShu, with the ATP-bound state being most favorable for DNA interaction.

**Table 2.**
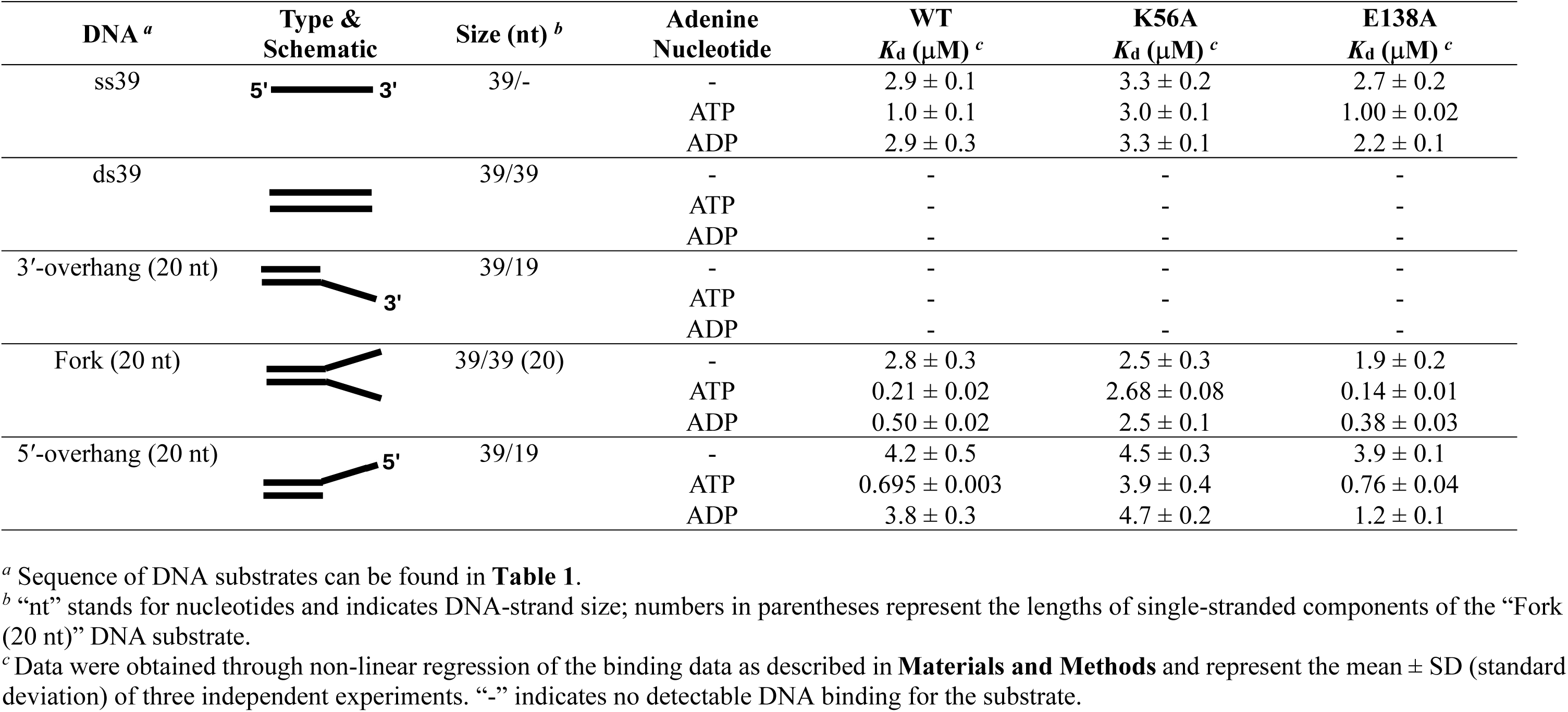
DNA-binding affinities of cShu and Walker-motif mutants (K56A, E138A) in the presence of ATP and ADP.

| DNA <sup>a</sup> | Type & Schematic | Size (nt) <sup>b</sup> | Adenine Nucleotide | WT $K_d$ (μM) <sup>c</sup> | K56A $K_d$ (μM) <sup>c</sup> | E138A $K_d$ (μM) <sup>c</sup> |
| --- | --- | --- | --- | --- | --- | --- |
| ss39 | 5' 3' | 39/- | - | 2.9 ± 0.1 | 3.3 ± 0.2 | 2.7 ± 0.2 |
| | | | ATP | 1.0 ± 0.1 | 3.0 ± 0.1 | 1.00 ± 0.02 |
| | | | ADP | 2.9 ± 0.3 | 3.3 ± 0.1 | 2.2 ± 0.1 |
| ds39 |  | 39/39 | - | - | - | - |
|  |  |  | ATP | - | - | - |
|  |  |  | ADP | - | - | - |
| 3'-overhang (20 nt) |  | 39/19 | - | - | - | - |
|  |  |  | ATP | - | - | - |
|  |  |  | ADP | - | - | - |
| Fork (20 nt) | | 39/39 (20) | - | 2.8 ± 0.3 | 2.5 ± 0.3 | 1.9 ± 0.2 |
| | | | ATP | 0.21 ± 0.02 | 2.68 ± 0.08 | 0.14 ± 0.01 |
| | | | ADP | 0.50 ± 0.02 | 2.5 ± 0.1 | 0.38 ± 0.03 |
| 5'-overhang (20 nt) | | 39/19 | - | 4.2 ± 0.5 | 4.5 ± 0.3 | 3.9 ± 0.1 |
| | | | ATP | 0.695 ± 0.003 | 3.9 ± 0.4 | 0.76 ± 0.04 |
| | | | ADP | 3.8 ± 0.3 | 4.7 ± 0.2 | 1.2 ± 0.1 |

<sup>a</sup> Sequence of DNA substrates can be found in **Table 1**.
<sup>b</sup> “nt” stands for nucleotides and indicates DNA-strand size; numbers in parentheses represent the lengths of single-stranded components of the “Fork (20 nt)” DNA substrate.
<sup>c</sup> Data were obtained through non-linear regression of the binding data as described in **Materials and Methods** and represent the mean ± SD (standard deviation) of three independent experiments. “-” indicates no detectable DNA binding for the substrate.

To validate these findings, we repeated the analysis using the Walker A (K56A) and Walker B (E138A) motif mutants of RFS1 in the cShu trimer (**Table 2**, **Figure S1G – S1I**). These key residue substitutions specifically abolish the ATP-binding (K56A) and ATP-hydrolysis (E138A) functions of RFS1-RIP1-SWS1, respectively. The mutants were purified (**Figure 1A & 1B**), and basic DNA-binding capabilities were verified by EMSAs (**Figure S1J**). As expected, the ATP-binding-deficient K56A mutant lost adenine nucleotide binding (**Figure 2A**) and failed to modulate DNA binding (**Table 2**). In contrast, the E138A mutant, which retains adenine nucleotide-binding ability (**Figure 2A**), maintained adenine nucleotide-dependent DNA-binding affinities comparable to those of the wild-type (WT) complex (**Table 2**). Thus, the mutant DNA-binding data confirm that ATP binding increases the DNA-binding specificity of cShu. Taken together, our results indicate that cShu preferentially binds DNA with an exposed 5′ end, requiring a sufficiently long ssDNA region for optimal interaction, with ssDNA/dsDNA junctions further contributing to binding. These DNA-binding preferences suggest a conserved substrate-recognition mechanism among Shu complex homologs, supporting a role for cShu in recognizing the lagging strand at stalled replication forks during HR^24,26,39^.

**Figure 2.**
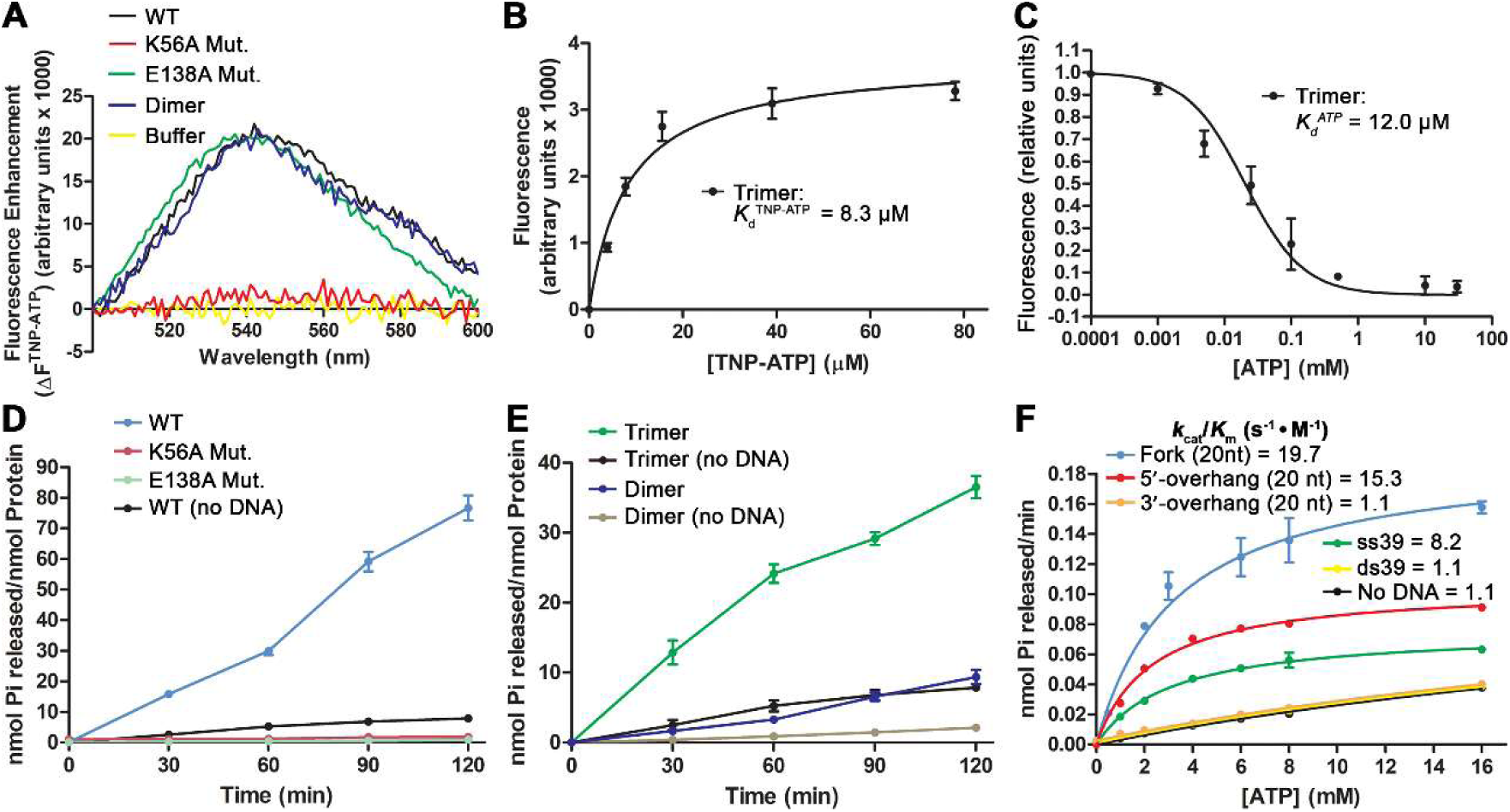
ATP binding and ATPase activity of trimeric cShu. (**A**) Fluorescence spectra of TNP-ATP (5 μM) with WT trimeric cShu, Walker-motif mutants (K56A, E138A), and the dimeric RFS1-RIP1 subcomplex (4 μM). (**B**) TNP-ATP saturation assay. Increasing concentrations of TNP-ATP were added to cShu (2 μM). Dissociation constants (*K*_d_^TNP-ATP^) were determined by non-linear curve fitting using a one-site binding model. (**C**) Titration of TNP-ATP (2.5 μM) fluorescence with ATP in the presence of cShu (2 μM). Data shown represent fluorescence units after subtraction of TNP-ATP background fluorescence, followed by normalization to the reading without ATP (relative units). Triplicate data were fitted using non-linear regression to a one-site competitive binding model. Dissociation constants (*K*_d_^ATP^) were derived from competitive binding data (see **Materials and Methods**). (**D**) ATP hydrolysis by WT cShu and Walker-motif mutants (1 μM each) with the dsDNA substrate, Fork (20 nt) (10 μM), over a 120-minute time course. (**E**) ATP hydrolysis by trimeric cShu and the dimeric RFS1-RIP1 subcomplex (1 μM each) with the ssDNA substrate, ss39 (10 μM), over a 120-minute time course. (**F**) ATPase activity (60-minute reactions) of cShu (1 μM) in the presence of ssDNA and dsDNA substrates with different end types (1 μM each), with increasing ATP concentrations. Data were fitted using the Michaelis–Menten model to determine kinetic parameters with the associated standard deviation from triplicate experiments (details found in **Table 3**). The number in parentheses following a given substrate name is the number of nucleotides in the single-stranded component(s) of the dsDNA substrates in **D** and **F**. Data in **D–F** represent the mean of three independent replicates, with error bars indicating the standard deviation from triplicate experiments.

**Table 3.** ATPase activity of trimeric cShu.

| DNA <sup>a</sup> | Type & Schematic | $K_m$ (mM) <sup>b</sup> | $V_{max}$<br>($\mu\text{M Pi} \cdot \text{min}^{-1}$) <sup>b</sup> | $k_{cat}$ ($\text{min}^{-1}$) <sup>c</sup> | $k_{cat}/K_m$<br>($\text{s}^{-1} \cdot \text{M}^{-1}$) |
| --- | --- | --- | --- | --- | --- |
| - | - | $39.2 \pm 0.1$ | $2.572 \pm 0.003$ | 2.6 | 1.1 |
| ss39 | 5'— 3' | $3.1 \pm 0.2$ | $1.53 \pm 0.04$ | 1.5 | 8.2 |
| ds39 | == | $35.2 \pm 0.1$ | $2.388 \pm 0.004$ | 2.4 | 1.1 |
| Fork (20 nt) | ==> | $3.3 \pm 0.1$ | $3.9 \pm 0.2$ | 3.9 | 19.7 |
| 3'-overhang (20 nt) | ==> 3' | $38.1 \pm 0.1$ | $2.534 \pm 0.004$ | 2.5 | 1.1 |
| 5'-overhang (20 nt) | ==< 5' | $2.3 \pm 0.2$ | $2.11 \pm 0.03$ | 2.1 | 15.3 |

<sup>a</sup> Sequence of DNA substrates can be found in **Table 1**.
<sup>b</sup> Data were obtained through non-linear regression of the ATPase data as described in **Materials and Methods** and represent the mean ± SD (standard deviation) of three independent experiments.
<sup>c</sup> $k_{cat} = V_{max}/[\text{cShu}]$ , where $[\text{cShu}] = 1 \mu\text{M}$ .

### cShu binds ATP through the Walker A domain of RFS1

RAD51 paralogs, adopting a RAD51-like fold, contain highly conserved Walker A (ATP-binding) and Walker B (ATP-hydrolysis) domains essential for regulating HR-associated processes^40–43^. As a RAD51 paralog complex, the RFS1-RIP1 dimer was previously shown to modulate RAD51 filaments in a manner strictly dependent on ATP binding^21^. To determine whether the trimeric cShu deviates from the dimeric RFS1-RIP1 subcomplex, we first analyzed the ATP binding of RFS1-RIP1-SWS1 and the Walker-motif mutants. We performed ATP-binding assays using a fluorescent ATP analog, 2′(3′)-O-(2,4,6-trinitrophenyl) adenosine 5′-triphosphate (TNP-ATP), to measure fluorescence intensity changes induced by the trimer variants and the purified RFS1-RIP1 dimer (**Figure S2A & S2B**). The WT cShu trimer increased the fluorescence intensity with a blue shift of the emission peak from 561 nm (TNP-ATP alone) to 542 nm (**black line in Figure 2A**), overlapping with that of the dimer (**blue line in Figure 2A**). These results confirm that both the cShu trimer and dimer bind TNP-ATP, sequestering the fluorescent ATP analog from the aqueous solution. Similar results were observed with the yeast Shu complex, in which the complete tetramer exhibited ATP binding comparable to the dimeric constituent, Csm2-Psy3^24^. The K56A (Walker A) mutant trimer completely lost the ability to bind TNP-ATP, with fluorescence changes reduced to baseline buffer levels (**Figure 2A**). In contrast, the E138A (Walker B) mutant trimer, which disrupts ATP hydrolysis without affecting binding, showed a fluorescence change and blue shift comparable to the WT trimer (**Figure 2A**). These results confirm that ATP binding in the *C*. *elegans* Shu complex is specifically conferred by the RFS1 subunit through the canonical Walker A motif, and that these point mutations did not introduce unintended functional disruptions to the complex.

Furthermore, the data indicate that the SWS1 subunit is not required for adenine nucleotide binding, consistent with the yeast Csm2-Psy3 dimer^24^.

With TNP-ATP binding confirmed, we determined the dissociation constant (*K*_d_^TNP-ATP^) of cShu for TNP-ATP to be 8.3 μM (**Figure 2B**). We further assessed the adenine nucleotide binding by performing competition assays using excess ATP and ADP (2.5 mM) against TNP-ATP (5 μM), which resulted in similar fluorescence reductions for both ATP and ADP. This indicates that ATP and ADP bind competitively to the same site without a strong preference for either adenine nucleotide (**Figure S2E**). Finally, we determined the dissociation constant for ATP (*K*_d_^ATP^) to be 12.0 μM (**Figure 2C**). As the intracellular ATP concentration in eukaryotic cells is ∼3 mM^44,45^, the *C*. *elegans* Shu complex is likely ATP-bound under physiological conditions, poised to perform HR-associated functions.

### cShu exhibits DNA-stimulated ATPase activity mediated by the Walker B domain of RFS1

The Shu complex homologs in yeast and humans have both been shown to exhibit ATPase activity^10,24,26^. The yeast Shu tetramer functions strictly as a DNA-dependent ATPase^24^, while the human Shu complex exhibits DNA-stimulated ATPase activity with weak basal hydrolysis in the absence of DNA^10^. The RFS1-RIP1 dimer was reported to exhibit barely detectable activity, reflecting an intrinsic deficiency due to the absence of the SWS1 subunit^12^. Since ATP hydrolysis is a conserved feature among Shu homologs, we expected that the cShu trimer, complete with the SWIM domain of SWS1, would hydrolyze ATP to fulfill the role of the complex as a RAD51 mediator.

We assessed the ATPase activity of cShu using malachite green-based ATP hydrolysis assays in the presence of the fork-shaped dsDNA substrate, Fork (20 nt). The cShu trimer exhibited markedly enhanced ATPase activity in the presence of the DNA substrate (**Figure 2D & 2E**). In contrast, the RFS1-RIP1 dimer only exhibited a DNA-stimulated activity close to the basal level of the trimer, substantially weaker than the DNA-stimulated activity of the complete trimer (**Figure 2E**). These observations indicate that the SWIM-containing subunit, SWS1, is required for normal ATPase activity of cShu, consistent with findings that conserved SWIM domains are indispensable for the ATPase activity in yeast and human Shu complexes^10,24,26^. As negative controls, both Walker-motif mutants (K56A and E138A in RFS1) completely lacked ATPase activity (**Figure 2D**). These results demonstrate that the *C*. *elegans* Shu complex exhibits intrinsic DNA-stimulated ATPase activity, conferred by the Walker A and B motifs within the RFS1 subunit.

To quantitatively analyze the ATPase activity and investigate substrate specificity, we determined the kinetic parameters in the presence of different DNA substrates (**Figure 2F**). The *k*_cat_ values ranged from 1.53 to 3.86 min^-1^, indicating a slow ATPase likely involved in regulatory functions (**Table 3**). The catalytic efficiencies (*k*_cat_/*K*_m_) for ss39, 5′-overhang, and Fork (20 nt) substrates were 8.2, 15.3, and 19.7 s^-1^ · M^-1^, respectively (**Table 3**). Notably, the catalytic efficiencies for the 5′-overhang and fork-shaped substrates, both featuring ssDNA/dsDNA junctions with an exposed 5′ end, were ∼15-fold and ∼20-fold higher than that in the absence of DNA, respectively. In contrast, the *k*_cat_/*K*_m_ values for the blunt-ended ds39 and the 3′-overhang substrates were equivalent to that of cShu in the absence of DNA (1.1 s^-1^ · M^-1^) (**Table 3**), consistent with the lack of detectable DNA binding to these substrates (**Table 1 & 2**). These results indicate that an exposed 5′ end at ssDNA/dsDNA junctions is critical for optimal ATPase activity of cShu. The preference for 5′-end-exposed ssDNA/dsDNA substrates in ATPase stimulation further suggests that cShu selectively recognizes specific DNA structures to regulate catalytic activity, a property also observed in the human Shu complex^26^. Similarly, the yeast Shu complex preferentially binds fork-shaped substrates with lesions on the lagging strand (5′-end-exposed arm)^39^. Thus, this strand-specific recognition mechanism is conserved and coupled to the preference for 5′-end-exposed DNA substrates observed across the Shu complexes.

### cShu alters RAD51 filament conformation and stability in an ATP-dependent manner

To assess the influence of the cShu trimer on RAD51 filaments, we examined filament accessibility using DNase I digestion protection assays. Poly(dT) ssDNA substrates were used to eliminate potential sequence bias. Using purified RAD51 (**Figure S3**), we conducted two-hour DNase I digestion assays on RAD51-ssDNA filaments in the presence of RFS1-RIP1-SWS1. As expected, RAD51-ssDNA filaments protected the DNA from DNase I digestion (**lane 6 vs. lanes 2 & 4 in Figure 3A & 3B**). Increasing concentrations of cShu sensitized the digestion of both 5′- and 3′-labeled RAD51 filaments in a concentration-dependent manner (**lanes 7–10 in Figure 3A & 3B**), indicating increased filament accessibility. The increased filament accessibility, regardless of fluorophore-labeling position, suggests that cShu induces conformational changes along the entire filament, consistent with the remodeling activity previously reported for the RFS1-RIP1 dimer^12,21^. Notably, this remodeling activity has been shown to be ATP-dependent in the RFS1-RIP1 dimer^21^, as well as in the yeast and human Shu homologs^24,26^. To determine whether filament remodeling by the complete cShu trimer also depends on ATP, we performed the assays using the Walker-motif trimer mutants, which are defective in either ATP binding (K56A) or ATP hydrolysis (E138A). Both mutants failed to sensitize RAD51 filaments to DNase I digestion (**lanes 9 & 10 in Figure 3C & 3D**), demonstrating that RAD51 filament remodeling is an intrinsic ATP-dependent activity of the *C*. *elegans* Shu complex.

**Figure 3.**
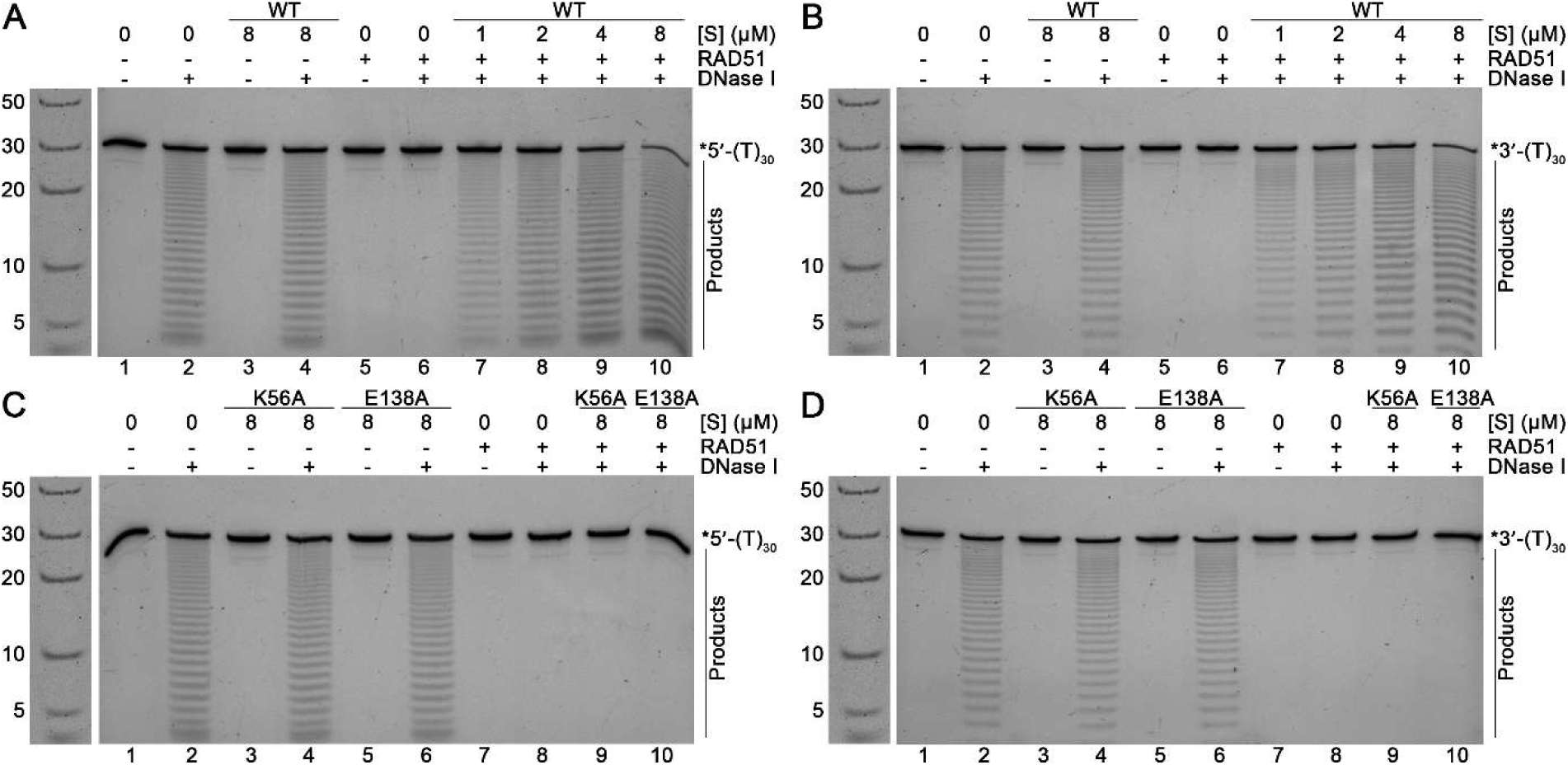
cShu enhances DNase I digestion on RAD51 filaments. (**A–B**) Two-hour DNase I digestion of RAD51-ssDNA filaments labeled on (**A, C**) the 5′ or (**B, D**) 3′ end with 6-FAM and incubated with/without WT cShu or Walker-motif mutants (K56A, E138A). Filaments (2 μM Rad51 on 0.25 μM poly(dT)_30_ ssDNA with 2 mM ATP) were incubated for 10 minutes and then mixed with increasing concentrations of WT cShu or mutants ([S]). Digestions were carried out using 2 Units DNase I at 37 °C. Reaction mixtures were resolved by 22.5% denaturing PAGE. DNA ladder with nucleotide sizes (in nt) is shown to the left of each gel and reused across all panels to indicate the sizes of the substrate and the corresponding degradation products. “*5′-(T)_30_” and “*3′-(T)_30_” stand for 5′- and 3′-labeled poly(dT)_30_ ssDNA, respectively.

Next, we used fluorescence-based assays to further investigate the RAD51-ssDNA filament conformational changes induced by the cShu trimer. In this assay, reductions in fluorescence correspond to increased solvent exposure of the fluorophore, indicating Shu complex-mediated conformational changes within the filament^12,21,24,26^. Fluorescence intensities of RAD51 filaments were measured at 10-second intervals over 60 seconds to capture early events in filament remodeling. The fluorescence of RAD51 filaments alone remained stable over the 60-second time frame, serving as a normalized baseline (**dashed black line in Figure 4A**). Upon addition of RFS1-RIP1-SWS1, the fluorescence intensity of the 5′-labeled RAD51-ssDNA filaments decreased by ∼35% within 10 seconds and by ∼45% at 60 seconds (**red line in Figure 4A**), whereas the 3′-labeled filaments showed no change over the same time frame (**pink line in Figure 4A**). This 5′-end polarity is consistent with previous findings for the RFS1-RIP1 dimer^21^. In comparison, the RFS1-RIP1 dimer induced only a ∼10% reduction at the 5′ end and none at the 3′ end (**Figure 4A**). The pronounced remodeling effect of the cShu trimer relative to the dimer correlates with the ATPase activities of the two complexes (**Figure 2E**), indicating that ATP hydrolysis is important for the action of cShu on RAD51 filaments. The 5′-end polarity of this effect indicates that filament remodeling is initiated specifically at the 5′ end, a conserved feature also observed in human and yeast Shu homologs^24,26^.

**Figure 4.**
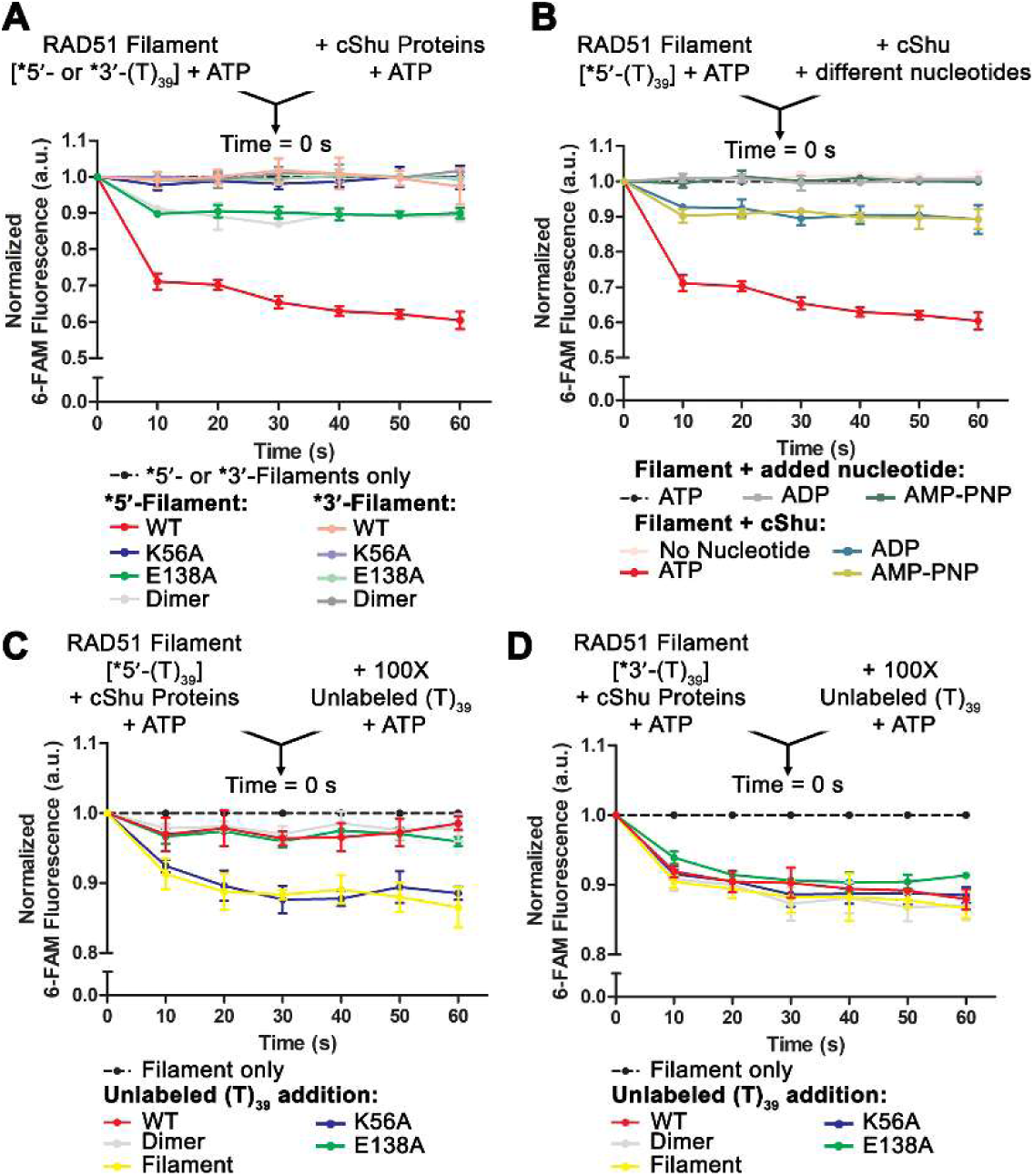
Conformational and stability changes of RAD51 filaments induced by trimeric cShu. (**A**) Fluorescence intensity profiles of RAD51-ssDNA filaments with/without cShu proteins over a 60-second time frame. Filaments (1 μM RAD51 on 15 nM 5′- or 3′-labeled poly(dT)_39_ ssDNA) were incubated for 10 minutes and then mixed with WT trimeric cShu, Walker-motif mutants (K56A, E138A), or the dimeric RFS1-RIP1 subcomplex (50 nM each), all in the presence of ATP (2 mM). (**B**) Fluorescence intensity profiles of RAD51-ssDNA filaments (pre-incubated with 2 mM ATP) mixed with cShu (pre-incubated with 2 mM of the indicated adenine nucleotide), over 60 seconds. (**C, D**) Fluorescence intensity profiles of RAD51-ssDNA filaments with/without WT trimeric cShu, Walker-motif mutants, or the dimeric RFS1-RIP1 subcomplex, mixed with a 100-fold excess of unlabeled ssDNA, over 60 seconds. Samples were excited at 490 nm, and emission readings were measured at 522 nm at equilibrium (20 °C). Filament-alone data were normalized as the baseline in arbitrary units corresponding to fluorescence intensities (dashed black lines). “*5′-(T)_39_” and “*3′-(T)_39_” stand for 5′- and 3′-labeled poly(dT)_39_ ssDNA, respectively.

To assess the adenine nucleotide dependence of filament remodeling, we performed the fluorescence assays with cShu preincubated with/without specific adenine nucleotide (ATP, ADP—the product of ATP hydrolysis—or AMP-PNP, a non-hydrolyzable ATP analog) before mixing with the filaments. Adenine nucleotide-free cShu did not induce detectable fluorescence changes (**Figure 4B**), similar to the ATP-binding-deficient Walker A mutant (**Figure 4A**), indicating cShu must be in an adenine nucleotide-bound state to act on filaments. Both ADP and AMP-PNP preincubations resulted in fluorescence reductions of ∼10%, significantly lower than with ATP (over 30%) (**Figure 4B**). These reductions were comparable to those of the ATP-hydrolysis-deficient Walker B mutant (which retains ATP binding) and the RFS1-RIP1 dimer (which exhibits barely detectable ATPase activity) (**Figure 4A**). These results demonstrate that ATP binding is required for cShu to induce a basal level of conformational changes in RAD51 filaments, whereas ATP hydrolysis is necessary to drive significant filament remodeling. Collectively, these findings highlight that SWS1 is critical for the ATPase activity of cShu and, in turn, for the capacity of the complex to remodel RAD51 filaments. The correlation between the ATPase activity and filament remodeling underscores a mechanistic coupling between the two processes. Consistent with the ATP-hydrolysis-dependent filament remodeling observed in yeast and human Shu homologs^24,26^, our data indicate that ATP hydrolysis by the *C*. *elegans* Shu complex is essential for driving conformational changes in RAD51 filaments.

Finally, we examined whether the trimeric cShu affects the stability of RAD51 filaments. We monitored filament fluorescence upon challenge with a 100-fold excess of unlabeled ssDNA over 60 seconds. The fluorescence reductions reflect the replacement of labeled ssDNA within the filaments by unlabeled ssDNA. In the absence of cShu, RAD51 filament fluorescence decreased by ∼10% and then remained stable (**yellow lines in Figure 4C & 4D**). Upon addition of cShu, the fluorescence reduction of 5′-labeled filaments was attenuated to ∼3%, with no significant effect on 3′-labeled filaments (**red lines in Figure 4C & 4D**). These results indicate that cShu suppresses DNA exchange at the 5′ end of RAD51 filaments, thereby stabilizing the filaments over the 60-second time frame, while the 3′ end remains unaffected. This preferential 5′-end stabilization by the trimer is consistent with our dimer controls (**Figure 4C & 4D**), as well as previous observations for the RFS1-RIP1 dimer^21^ and human and yeast Shu homologs^24,26^. The Walker-motif mutants were used to distinguish the contributions of ATP binding and hydrolysis to filament stabilization. The ATP-binding mutant (K56A) completely lost the ability to stabilize RAD51 filaments (**blue line in Figure 4C**). In contrast, the ATP-hydrolysis mutant (E138A), which retains ATP binding, stabilized filaments at the 5′ end as effectively as the WT trimer (**green and red lines in Figure 4C**). These results demonstrate that ATP binding, rather than hydrolysis, is essential for RFS1-RIP1-SWS1-mediated stabilization of RAD51 filaments. Overall, our findings show that the *C*. *elegans* Shu complex stabilizes RAD51 filaments in an ATP-binding-dependent manner, while ATP hydrolysis drives conformational remodeling. The intrinsic ability of cShu to modulate RAD51 filaments highlights the essential role of the complex as a RAD51 mediator.

## Discussion

Biochemical characterization of the *C*. *elegans* Shu complex provides key insights into the role of the complex in modulating RAD51 filaments and regulating HR during the DNA damage tolerance response. Quantitative DNA-binding analysis (*K*_d_) revealed that the cShu trimer preferentially binds DNA substrates with an exposed 5′ end, particularly the fork-shaped dsDNA with 20-nt arms in the presence of ATP. Notably, our experiments were conducted using undamaged DNA substrates, indicating that the observed binding preferences are intrinsic to RFS1-RIP1-SWS1. Similar preferences have been reported for the yeast and human Shu homologs^26,39^, aligning with the role of the yeast Shu complex in lesion bypass on lagging strands at replication forks^39^. These findings suggest that substrate recognition is a conserved feature of Shu complexes across eukaryotic species. The selective binding of exposed 5′ ends over 3′ ends of DNA further supports a lagging-strand-specific interaction at stalled replication forks (**Figure 5**). Nevertheless, the DNA-binding affinities of Shu homologs have been shown to be lower than the affinities of species-specific Rad51 proteins^24,26,46,47^. Given that RAD51 is likely more abundant than the Shu complex in cells, it is expected to dominate DNA binding, while cShu serves primarily as a regulatory factor to modulate filament dynamics.

**Figure 5.**
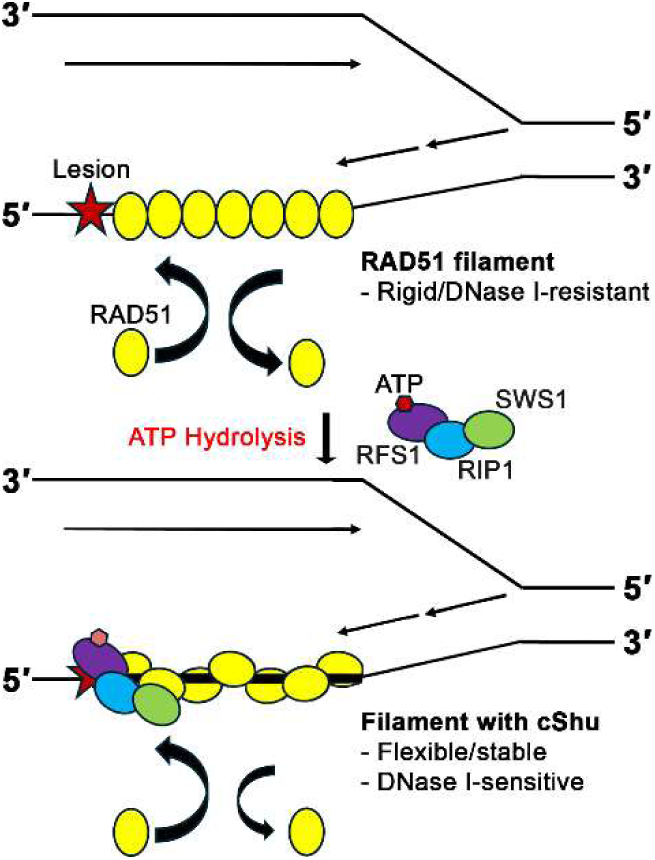
Molecular model for the action of cShu on RAD51-ssDNA filaments. Proposed mechanism of cShu as a DNA-stimulated ATPase in homologous recombination. cShu binds to the 5′ end of RAD51 filaments at sites containing DNA lesions, such as abasic lesions, where it stabilizes the filament in an ATP-binding-dependent manner. The remodeling of RAD51 filament conformation is dependent on the ATPase activity of cShu, rendering the filament accessible and “open” for HR-associated processes.

Our study demonstrates that the RFS1 subunit contains both Walker A and B motifs, which confer ATP binding and hydrolysis to the *C*. *elegans* Shu complex. Importantly, our work shows that the SWS1 subunit is indispensable for functional ATP hydrolysis. Disruption of the SWIM domain impairs eukaryotic DNA damage tolerance^14^, consistent with a critical role in the ATPase activity of cShu and the yeast and human homologs^10,24^. These findings underscore the importance of ATPase activity and the functional integrity of this activity for the role of the Shu complex in HR-associated error-free DNA lesion bypass.

Building on previous studies of the RFS1-RIP1 dimer^12,21^, we found that ATP binding and hydrolysis in the cShu trimer serve distinct roles in modulating pre-assembled RAD51 filaments (**Figure 5**). The trimer requires ATP binding to induce a limited conformational change in RAD51 filaments, whereas ATP hydrolysis drives additional remodeling beyond these basal changes. Using site-specific Walker-motif mutants, we further determined that filament stabilization by cShu depends specifically on ATP binding. Distinct roles for ATP binding and hydrolysis in RAD51 filament modulation were also observed in the human Shu complex^26^. Overall, the SWS1 subunit is critical for the ATPase activity of RFS1-RIP1-SWS1, facilitating filament conformational changes that initiate at the 5′ end and propagate toward the 3′ end. This function likely underlies the role of the complex as a RAD51 mediator that promotes downstream HR-associated DNA damage response. In closing, our work highlights the essential role of the DNA-stimulated ATPase activity of the *C*. *elegans* Shu complex in regulating RAD51 filament properties, a conserved feature among Shu complex homologs that supports error-free DNA lesion bypass.

## Data Availability

Source data are available in **Supporting Information** or upon request.

## Acknowledgements

We thank Dr. Simon J. Boulton for providing the plasmid of the *C*. *elegans* RAD51. The work was supported by The Natural Sciences and Engineering Research Council of Canada (through a Discovery grant (RGPIN/06165-2019) to H.L.).

## Abbreviations

HR, homologous recombination ssDNA, single-stranded DNA dsDNA, double-stranded DNA

EMSA, electrophoretic mobility shift assay PAGE, polyacrylamide gel electrophoresis FPA, fluorescence polarization assay TNP-ATP, 2′(3′)-O-(2,4,6-trinitrophenyl) adenosine 5′-triphosphate WT, wild type

## Author Contributions

**S. S. H. C.:** Conceptualization, Methodology, Validation, Investigation, Data analysis, Writing, & Editing.

**G. X.:** Methodology, Investigation.

**H. L.:** Conceptualization, Data analysis, Writing, Review & Editing, Project administration, Funding acquisition.

## Competing Interest Statement

No author has an actual or perceived conflict of interest with the contents of this article.

## Notes

### Competing Interest Statement

The authors have declared no competing interest.

### Summary of Updates

Spelling, grammatical and punctuation errors corrected; "Figure 6" reference in Discussion Section corrected to "Figure 5"

## References

1. Marteijn, J. A., Lans, H., Vermeulen, W. & Hoeijmakers, J. H. J. Understanding nucleotide excision repair and its roles in cancer and ageing. Nat. Rev. Mol. Cell Biol. 15, 465–481 (2014).

2. van Gent, D. C., Hoeijmakers, J. H. & Kanaar, R. Chromosomal stability and the DNA double-stranded break connection. Nat. Rev. Genet. 2, 196–206 (2001).

3. Branzei, D. & Foiani, M. Regulation of DNA repair throughout the cell cycle. Nat. Rev. Mol. Cell Biol. 9, 297–308 (2008).

4. Moynahan, M. E. & Jasin, M. Mitotic homologous recombination maintains genomic stability and suppresses tumorigenesis. Nat. Rev. Mol. Cell Biol. 11, 196–207 (2010).

5. Kowalczykowski, S. C. An Overview of the Molecular Mechanisms of Recombinational DNA Repair. Cold Spring Harb. Perspect. Biol. 7, a016410 (2015).

6. Carver, A. & Zhang, X. Rad51 filament dynamics and its antagonistic modulators. Semin. Cell Dev. Biol. 113, 3–13 (2021).

7. San Filippo, J., Sung, P. & Klein, H. Mechanism of eukaryotic homologous recombination. Annu. Rev. Biochem. 77, 229–257 (2008).

8. Sung, P. & Klein, H. Mechanism of homologous recombination: mediators and helicases take on regulatory functions. Nat. Rev. Mol. Cell Biol. 7, 739–750 (2006).

9. Prakash, R., Zhang, Y., Feng, W. & Jasin, M. Homologous recombination and human health: the roles of BRCA1, BRCA2, and associated proteins. Cold Spring Harb. Perspect. Biol. 7, a016600 (2015).

10. Liu, T., Wan, L., Wu, Y., Chen, J. & Huang, J. hSWS1·SWSAP1 Is an Evolutionarily Conserved Complex Required for Efficient Homologous Recombination Repair. J. Biol. Chem. 286, 41758–41766 (2011).

11. Martín, V. et al. Sws1 is a conserved regulator of homologous recombination in eukaryotic cells. EMBO J. 25, 2564–2574 (2006).

12. Taylor, M. R. G. et al. Rad51 Paralogs Remodel Pre-synaptic Rad51 Filaments to Stimulate Homologous Recombination. Cell 162, 271–286 (2015).

13. McClendon, T. B., Sullivan, M. R., Bernstein, K. A. & Yanowitz, J. L. Promotion of Homologous Recombination by SWS-1 in Complex with RAD-51 Paralogs in Caenorhabditis elegans. Genetics 203, 133–145 (2016).

14. Godin, S. K. et al. Evolutionary and Functional Analysis of the Invariant SWIM Domain in the Conserved Shu2/SWS1 Protein Family from Saccharomyces cerevisiae to Homo sapiens. Genetics 199, 1023–1033 (2015).

15. Bernstein, K. A. et al. The Shu complex, which contains Rad51 paralogues, promotes DNA repair through inhibition of the Srs2 anti-recombinase. Mol. Biol. Cell 22, 1599–1607 (2011).

16. Shor, E., Weinstein, J. & Rothstein, R. A Genetic Screen for top3 Suppressors in Saccharomyces cerevisiae Identifies SHU1, SHU2, PSY3 and CSM2: Four Genes Involved in Error-Free DNA Repair. Genetics 169, 1275 (2005).

17. Matsuzaki, K., Kondo, S., Ishikawa, T. & Shinohara, A. Human RAD51 paralogue SWSAP1 fosters RAD51 filament by regulating the anti-recombinase FIGNL1 AAA+ ATPase. Nat. Commun. 10, 1–15 (2019).

18. Keeney, S., Giroux, C. N. & Kleckner, N. Meiosis-specific DNA double-strand breaks are catalyzed by Spo11, a member of a widely conserved protein family. Cell 88, 375–384 (1997).

19. Cole, F., Keeney, S. & Jasin, M. Evolutionary conservation of meiotic DSB proteins: more than just Spo11. Genes Dev. 24, 1201–1207 (2010).

20. Hodgkin, J., Horvitz, H. R. & Brenner, S. Nondisjunction Mutants of the Nematode CAENORHABDITIS ELEGANS. Genetics 91, 67–94 (1979).

21. Taylor, M. R. G. et al. A Polar and Nucleotide-Dependent Mechanism of Action for RAD51 Paralogs in RAD51 Filament Remodeling. Mol. Cell 64, 926–939 (2016).

22. Špírek, M., Taylor, M. R. G., Belan, O., Boulton, S. J. & Krejci, L. Nucleotide proofreading functions by nematode RAD51 paralogs facilitate optimal RAD51 filament function. Nat. Commun. 12, 5545 (2021).

23. Belan, O. et al. Single-molecule analysis reveals cooperative stimulation of Rad51 filament nucleation and growth by mediator proteins. Mol. Cell 81, 1058–1073.e7 (2021).

24. Chu, S. S. H., Xing, G., Jha, V. K. & Ling, H. The Shu complex is an ATPase that regulates Rad51 filaments during homologous recombination in the DNA damage response. DNA Repair 145, 103792 (2025).

25. Hengel, S. R. et al. The human Shu complex promotes RAD51 activity by modulating RPA dynamics on ssDNA. Nat. Commun. 15, 7197 (2024).

26. Chu, S. S. H., Xing, G. & Ling, H. The role of human Shu complex in ATP-dependent regulation of RAD51 filaments during homologous recombination-associated DNA damage response. J. Biol. Chem. 301, 110212 (2025).

27. Klock, H. E., Koesema, E. J., Knuth, M. W. & Lesley, S. A. Combining the polymerase incomplete primer extension method for cloning and mutagenesis with microscreening to accelerate structural genomics efforts. Proteins 71, 982–994 (2008).

28. Scholz, J. & Suppmann, S. A new single-step protocol for rapid baculovirus-driven protein production in insect cells. BMC Biotechnol. 17, 83 (2017).

29. Tao, Y. et al. Structural Analysis of Shu Proteins Reveals a DNA Binding Role Essential for Resisting Damage. J. Biol. Chem. 287, 20231–20239 (2012).

30. Godin, S. et al. The Shu complex interacts with Rad51 through the Rad51 paralogues Rad55–Rad57 to mediate error-free recombination. Nucleic Acids Res. 41, 4525–4534 (2013).

31. LaConte, L. E. W., Srivastava, S. & Mukherjee, K. Probing Protein Kinase-ATP Interactions Using a Fluorescent ATP Analog. Methods Mol. Biol. Clifton NJ 1647, 171–183 (2017).

32. Stewart, R. C., VanBruggen, R., Ellefson, D. D. & Wolfe, A. J. TNP-ATP and TNP-ADP as Probes of the Nucleotide Binding Site of CheA, the Histidine Protein Kinase in the Chemotaxis Signal Transduction Pathway of Escherichia coli. Biochemistry 37, 12269–12279 (1998).

33. Lee, J. H., Garboczi, D. N., Thomas, P. J. & Pedersen, P. L. Mitochondrial ATP synthase. cDNA cloning, amino acid sequence, overexpression, and properties of the rat liver alpha subunit. J. Biol. Chem. 265, 4664–4669 (1990).

34. Ward, L. D. [22] Measurement of ligand binding to proteins by fluorescence spectroscopy. In Methods in Enzymology vol. 117 400–414 (Academic Press, 1985).

35. Lakiwicz, J. R. Quenching of Fluorescence. in Principles of Fluorescence Spectroscopy 238–266 (Plenum Press, New York, NY, USA, 1999).

36. Chan, K.-M., Delfert, D. & Junger, K. D. A direct colorimetric assay for Ca2+-stimulated ATPase activity. Anal. Biochem. 157, 375–380 (1986).

37. Rowlands, M. G. et al. High-throughput screening assay for inhibitors of heat-shock protein 90 ATPase activity. Anal. Biochem. 327, 176–183 (2004).

38. Godin, S. et al. The Shu complex interacts with Rad51 through the Rad51 paralogues Rad55-Rad57 to mediate error-free recombination. Nucleic Acids Res. 41, 4525–4534 (2013).

39. Rosenbaum, J. C. et al. The Rad51 paralogs facilitate a novel DNA strand specific damage tolerance pathway. Nat. Commun. 10, 3515 (2019).

40. Thacker, J. A surfeit of RAD51-like genes? Trends Genet. TIG 15, 166–168 (1999).

41. Thacker, J. The RAD51 gene family, genetic instability and cancer. Cancer Lett. 219, 125–135 (2005).

42. Thompson, L. H. & Schild, D. The contribution of homologous recombination in preserving genome integrity in mammalian cells. Biochimie 81, 87–105 (1999).

43. Thompson, L. H. & Schild, D. Homologous recombinational repair of DNA ensures mammalian chromosome stability. Mutat. Res. 477, 131–153 (2001).

44. Imamura, H. et al. Visualization of ATP levels inside single living cells with fluorescence resonance energy transfer-based genetically encoded indicators. Proc. Natl. Acad. Sci. U. S. A. 106, 15651–15656 (2009).

45. Takaine, M., Ueno, M., Kitamura, K., Imamura, H. & Yoshida, S. Reliable imaging of ATP in living budding and fission yeast. J. Cell Sci. 132, jcs230649 (2019).

46. Chabot, T. et al. New Phosphorylation Sites of Rad51 by c-Met Modulates Presynaptic Filament Stability. Cancers 11, 413 (2019).

47. Tombline, G., Shim, K.-S. & Fishel, R. Biochemical Characterization of the Human RAD51 Protein. J. Biol. Chem. 277, 14426–14433 (2002).

