## Supporting Information for "The trimeric Shu complex in *C. elegans* is an ATPase that remodels RAD51 filaments in the homologous recombination-associated DNA damage response"

**This PDF file includes:**

Figures S1 to S3

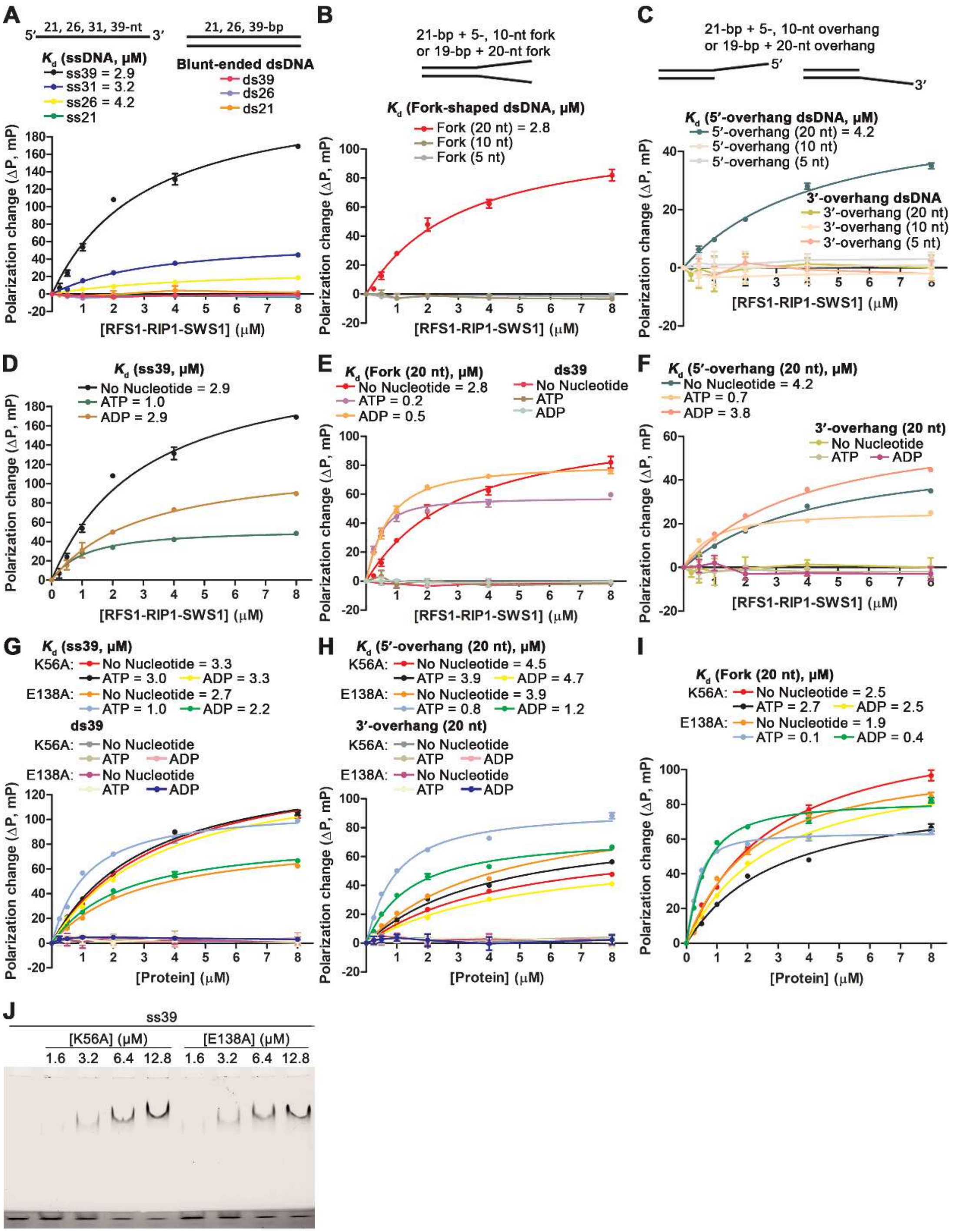


**Figure S1. DNA binding of trimeric *C. elegans* Shu complex (cShu) proteins.**

(**A–C**) Fluorescence polarization assay (FPA) for DNA binding of WT cShu with different ssDNA and dsDNA substrates (5 nM). (**D–I**) FPA for DNA binding of (**D–F**) WT cShu and (**G–I**) Walker-motif mutants (K56A and E138A in RFS1) with different DNA substrates (5 nM), in the presence of adenine nucleotides (2 mM). Dissociation constants ($K_d$) in **A–I** were determined by non-linear curve fitting to a one-site binding model (details in **Materials and Methods**). Data represent the mean of three independent replicates, with error bars indicating the standard deviation from triplicate experiments. The number in parentheses following a given substrate name is the number of nucleotides in the single-stranded component(s) of the dsDNA substrates in **B, C, E, F, H & I**. (**J**) Electrophoretic mobility shift assay (EMSA) for mutant cShu-DNA interactions. Increasing concentrations of Walker-motif mutants (K56A, E138A) were mixed with 0.05 μM ss39 DNA substrate. Protein-DNA mixtures were resolved by two-layer PAGE gels: 5% native polyacrylamide at the top and 15% at the bottom (dark layer). Assays were performed in triplicate with comparable results.

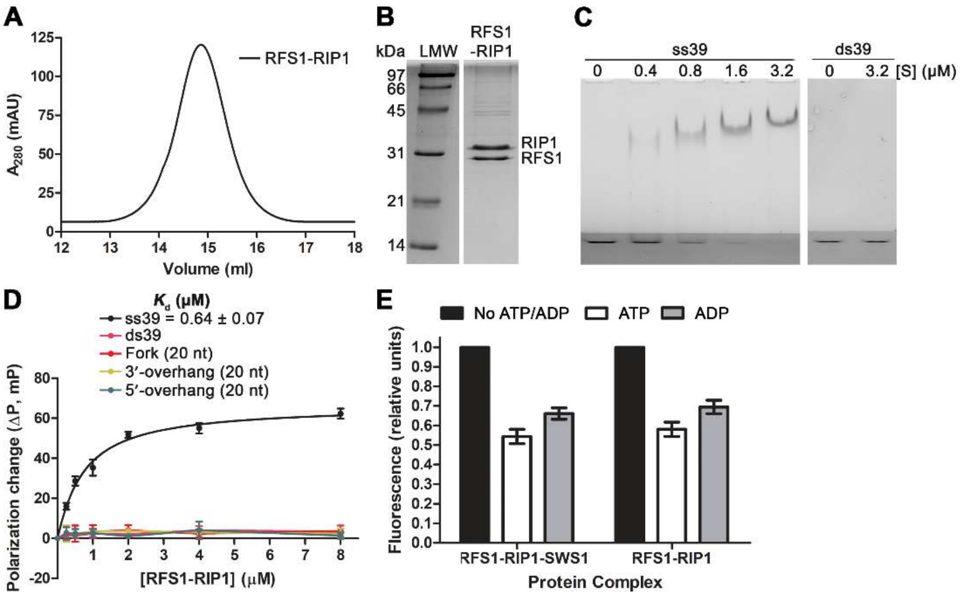


**Figure S2. Purified dimeric RFS1-RIP1 subcomplex and characterization of~~its~~ DNA and adenine nucleotide binding.**

(**A**) Gel filtration and (**B**) SDS-PAGE of the purified dimeric RFS1-RIP1 subcomplex. “LMW” stands for low-molecular-weight protein ladder. (**C**) Electrophoretic mobility shift assay (EMSA) for RFS1-RIP1-DNA interactions. Different concentrations of RFS1-RIP1 ([S]) were mixed with 0.05 μM DNA substrates (ss39 and ds39). Protein-DNA mixtures were resolved by two-layer

PAGE gels: 5% native polyacrylamide at the top and 15% at the bottom (dark layer). Assays were performed in triplicate with comparable results. (**D**) Fluorescence polarization assay (FPA) for DNA-binding of the dimeric RFS1-RIP1 subcomplex with different DNA substrates (5 nM). Dissociation constants ($K_d$) were determined by non-linear curve fitting to a one-site binding model (details in **Materials and Methods**). Data represent the mean of three independent replicates, with error bars indicating the standard deviation from triplicate experiments. (**E**) Competition assays of TNP-ATP (5 μM) bound to trimeric cShu or the dimeric RFS1-RIP1 subcomplex (4 μM) with ATP or ADP (2.5 mM each). Fluorescence units were normalized to the initial readings without excess ATP or ADP.

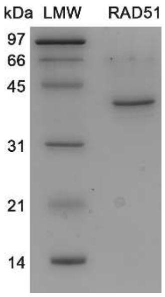


**Figure S3. Purified *C. elegans* RAD51 protein.**

SDS-PAGE analysis of the purified *C. elegans* RAD51. “LMW” stands for low-molecular-weight protein ladder.
